# A study of genetic variants associated with skin traits in the Vietnamese population

**DOI:** 10.1101/2023.09.06.556474

**Authors:** Tham Hong Hoang, Duc Minh Vu, Giang Minh Vu, Thien Khac Nguyen, Nguyet Minh Do, Vinh Chi Duong, Thang Luong Pham, Mai Hoang Tran, Ly Thi Khanh Nguyen, Han Thi Tuong Han, Thuy Thu Can, Thai Hong Pham, Tho Duc Pham, Thanh Hong Nguyen, Huy Phuoc Do, Nam S. Vo, Xuan-Hung Nguyen

## Abstract

**Background:** Most skin-related traits have been studied from Caucasian genetic background. A comprehensive study on skin-associated genetic effects on under-represented populations like Vietnam is needed to fill the gaps in the field.

**Objectives:** To develop a computational pipeline to predict the effect of genetic factors on skin traits using public data (GWAS catalogs and whole genome sequencing (WGS) data of 1000 genomes project-1KGP) and in-house Vietnamese data (WGS and genotyping by SNP array). By using this information we may have a better understanding of the susceptibility of Vietnamese people.

**Methods:** Vietnamese cohorts of whole genome sequencing (WGS) of 1008 healthy individuals for the reference and 96 genotyping samples (which do not have any skin cutaneous issues) by Infinium Asian Screening Array-24 v1.0 BeadChip were employed to predict skin-associated genetic variants of 25 skin-related and micronutrients requirement traits in population analysis and correlation analysis. Simultaneously, we compared the landscape of cutaneous issues of Vietnamese people with other populations by assessing their genetic profiles.

**Results:** The skin-related genetic profile of Vietnamese cohorts is similar at most with East Asian (JPT: Fst=0.036, CHB: Fst=0.031, CHS: Fst=0.027, CDX: Fst=0.025) in the population study. In addition, we identified pairs of skin traits being at high risk of frequent co-occurrence (such as skin aging and wrinkles (r = 0.45, p =1.50e-5) or collagen degradation and moisturizing (r = 0.35, p = 1.1e-3).

**Conclusion:** This is the first investigation in Vietnam to explore genetic variants of facial skin. These findings could improve inadequate skin-related genetic diversity in the currently published database.

## Introduction

As the largest organ in the human body, the skin acts as a barrier to regulate body temperature and prevent detrimental impacts from the external environment. Skin disorder is a common human health issue that severely weakens skin functions and hamper attractiveness [1]. Therefore, dermatological conditions are of immense concern for some individuals, especially females. As a result, there are increasing efforts to tackle these issues and improve skin health. One of the approaches is to study genetic factors affecting skin functions by providing in-depth understanding at genetic factors needed to construct proper prevention and treatment strategies.

Many exogenous factors, such as overall health, lifestyle, daily diets, and other environmental factors could influence skin condition. Micronutrient deficiencies could also be a potential causative factor leading to cutaneous abnormalities. The most well- documented cutaneous condition-related micronutrients are B, C, and several fat-soluble vitamins, such as A, E, D, and K [2]. Moreover, several studies have confirmed the essential roles of genetics in developing some skin-related phenotypes and the association of various genetic variants with skin pigmentation, freckles, hairiness, and excessive sweating in certain populations [3-7]. Recently, the advent of the Human Genome Project, genome-wide association studies (GWAS), and next-generation sequencing (NGS) enable experts to identify candidate loci in the genes underlying skin-related traits and skin disorders. The act of investigating the frequency and annotations of genetic variants on these loci in populations may shed light on specific skin disorders for each country as well as the difference in specific skin-related issues between populations.

Even though Asian populations make up over 40% of the global population and comprise a significant genetic diversity, however they are still underrepresented in current genomic studies. Up to now, skin genetics have been studied primarily on populations with European ancestry. The first Genome-wide association study (GWAS) of skin pigmentation included 1,043 individuals from 51 European ancestry populations who have identified major loci and specific polymorphisms affecting human skin color [7]. In 2018, a UK biobank-based GWAS was conducted to analyze a broad set of 23 highly inheritable traits, including Skin, Vitamin D levels, tanning, and sunburn [5]. Aside from Europeans, several studies have been done on Asian populations, such as Han Chinese, Indian, Japanese, and Korean [3, 4, 6, 8]. A large-scale GWAS of various skin phenotypes identified several novel skin-spot traits-associated signals neighbored *AKAP1/MSI2, BNC2, HSPA12A, PPARGC1B, RAB11FIP2*, and double-edged eyelid-related signal around *EMX2* in Japanese female population [3]. In Korea, a GWAS analysis in 17,019 women revealed several genomic loci significantly associated with facial pigmented spots (e.g., *BNC2, PPARGC1B, MC1R, MFSD12*) [6]. These insights helped dermatologic researchers better understand the genetic underpinnings of skin-related phenotypic variation in Asian populations.

There are several ways to estimate an individual’s genetic contribution to a phenotype. The two most popular methods are Polygenic Risk Score (PRS) [9] and GRS-RAC [10] (also called Count SNP Score or Top SNP Score). PRS estimates how the collection of one’s variants affects their risk for certain traits or diseases by using DNA information and data derived from large-scale genomic studies [9]. In contrast, the Top SNP Score only considers the genomic variants significantly associated with phenotypes. Therefore, Top SNP Score is much simpler than PRS but requires tremendous work for database curation [10]. However, both these approaches allow us to calculate the health condition-associated risks to guide healthcare decisions.

In the present study, we aimed to explore the unique pattern of the cutaneous problems in Vietnam by investigating the correlation among genetic and non-genetic factors (collected by customized questionnaire) associated with various traits, including skin-related and micronutrient requirements. In addition, we compared the landscape of cutaneous issues based on associated variants profiles in Vietnamese people with other populations worldwide to explain the inter-population variability in dermatological problems and provide understanding to improve dermatological healthiness.

## Materials and methods

### Study cohorts

i. Microarray dataset: This study included 96 Vietnamese adults with no remarkable medical history related to skin. After being consulted with dermatologists about the process and the objectives of the research, participants did a skin examination with a dermatoscope to assess dermatological issues (The assessment results were presented in **Table 1**). Then, clinicians interpreted the results and diagnosed the condition of the participant’s skin. All participants did a survey on other skin-related information. All participants provided consent to participate in this study using written informed consent forms. These samples were processed with the Infinium Asian Screening Array-24 v1.0 (ASA) BeadChip.
ii. 1000 Vietnamese Genomes (VN1K) dataset: We used whole genome sequencing (WGS) data of 1008 unrelated individuals from the 1000 Vietnamese Genomes Project (VN1K) by Vingroup Big Data Institute to investigate population-specific skin-related risks. In the VN1K study, subjects provided informed consent, and the study was approved by the Vinmec International Hospital Institutional Review Board with number 543/2019/QĐ-VMEC. The characteristics of 1008 individuals can be found on the official website of VN1K at https://genome.vinbigdata.org/.
iii. 1000 genome project phase 3 (1KGP3) dataset: We also used WGS data of 2504 unrelated individuals of five super populations of African (AFR), Ad Mixed American (AMR), East Asian (EAS), European (EUR), South Asian (SAS) for comparison with the VN1K population. This 1KGP3 dataset has its own consent form.

### DNA extraction, purification, and genotyping

DNA was extracted and purified using the MagMAX DNA Multi-Sample Ultra 2.0 Kit (ThermoFisher Scientific, USA) according to the manufacturer’s instructions. After the single-base extension, the ASA chip was processed and scanned by the Illumina iScan System. The ASA chip is designed with more than 660,000 variants for medical research and related disease studies in East Asian populations [11]. 94/96 samples have more than 98% calling rate. The .IDAT files (the format obtained from the Iscan machine for the ASA chip) were converted to .vcf, yielding 656,891 variants. Of these, 623,886 variants are located on autosomes. The research team used Shapeit4 [12] and Minimac4 [13] software to add imputation points with the genomes of 1000 Vietnamese Genomes people, removing bad variants with impute R2 < 0.3. Some criteria for selecting samples and variants: (i) samples: proportion of missing variants < 0.05 excluding blood-related samples, (ii) variants: missing proportions of variants to total samples < 0.05, variants with p-value < 10^−6^ following the Hardy-Weinberg principle, and with MAF (minor allele frequency) MAF < 0.01 were eliminated.

**Table 1.** Mean of VISIA assessment results.

| **Trait** | **Angle** | **Percentile** | **Feature count** | **Score** |
| --- | --- | --- | --- | --- |
| Spots | Left | 73.88 | 110.2 | 28.03 |
|  | Right | 75.95 | 108.3 | 26.95 |
|  | Front | 69.21 | 145.15 | 32.21 |
| Wrinkles | Left | 40.04 | 103.9 | 28.108 |
|  | Right | 42.23 | 107.4 | 27.377 |
|  | Front | 50.844 | 29.37 | 11.536 |
| Texture | Left | 46.23 | 1603 | 11.883 |
|  | Right | 43.3 | 1660 | 12.568 |
|  | Front | 43.211 | 2550.67 | 49.097 |
| Pores | Left | 51.98 | 552.1 | 14.953 |
|  | Right | 51.91 | 556.8 | 15.121 |
|  | Front | 48.065 | 962.54 | 52.441 |
| UV spots | Left | 87.98 | 260.4 | 16.338 |
|  | Right | 86.33 | 261 | 16.661 |
|  | Front | 88.35 | 369.514 | 17.738 |
| Brown spots | Left | 63.17 | 89.17 | 17.005 |
|  | Right | 63.56 | 85.1 | 16.805 |
|  | Front | 64.92 | 120 | 17.39 |
| Red areas | Left | 66.81 | 12.62 | 11.416 |
|  | Right | 65.25 | 12.73 | 11.616 |
|  | Front | 70.178 | 15.22 | 10.823 |
| Porphyrins | Left | 41.4 | 1588 | 17.421 |
|  | Right | 43.52 | 1564 | 17.104 |
|  | Front | 45.98 | 2563.26 | 71.155 |

### Evaluation of 25 common skin-related traits

Skin disorders vary greatly in symptoms and severity. They can be temporary or permanent and directly influence the quality of life. There are many different types of skin abnormalities. We investigated the 25 most common features: freckles, tanning response, the ability of UV protection, skin aging, elasticity, antioxidant capacity, and skin-related micronutrient requirement. Some of these present relationships with others; one issue could be the cause or the consequence of another. Generally, they could be categorized into four main groups: pigmentation, dermatitis, nutrition, and aging (Supplementary file, **Figure S1**).

### GWAS and SNP dataset curation for analysis

GWAS and other genetic association studies for skin features were collected from GWAS Catalog and Pubmed. In GWAS Catalog, the search strategy was based on the keywords and their synonyms for 25 skin-related traits (i.e., “Freckle,” “Tanning response,” “UV protection,” “Skin sensitivity,” “Inflammatory cytokines,” “Skin aging,” “Elasticity,” “Antioxidant,” “Collagen,” “Stretch mark,” “Wrinkle,” “Glycation,” “Moisturizing,” “Acne,” “Eyelid,” “Vitamin A need’, “Vitamin B2 need”, “Vitamin B6 need”, “Vitamin B9 need”, “Vitamin B12 need”, “Vitamin C need,” “Vitamin D need,” “Vitamin K need,” “Omega need”). For searching studies from the Pubmed platform, we also used the aforementioned search terms combined with others representing genetic association studies, such as “Genome-wide association studies,” “genetic association,” “genomic study,” or “GWAS.”

Selected studies must report information regarding genetic variations associated with skin-related traits. We will observe summary data for every SNP that significantly modulates the risk of cutaneous issues. Data extraction form includes genetic variation information (SNP, gene, location), participant information (quantity, ethnicity, mean age, gender proportion, etc.), affected skin traits, and statistical data (effect-size (beta) and p-value of the associations). Identified variants were compared with other sets of SNPs from several vendors on the market, such as OmeCare (U.S.A.) and BioEasy (Malaysia), to get high confidence in consensus SNPs. The list of identified SNP overlapped with the Vietnamese Genetic Variation Database from VN1K to establish a comprehensive variant dataset related to dermatology.

### Questionnaire forms

The questionnaire was constructed to build holistic metadata of all participants. All 96 participants in the microarray dataset would fulfill the questionnaire regarding demographics (gender, age), body mass index (BMI), lifestyle (stress, frequency of exercise, sunlight exposure levels), hypersensitivity history, alcohol used, smoke exposure (active/passive), medications, nutrition (sweet eating habits, fruit-eating habits), skin problems in family members, and information about skin care (approaches, intensity, skin satisfaction levels). The characteristics and information retrieved from the questionnaire form were summarized in **Table 2**.

**Table 2.**
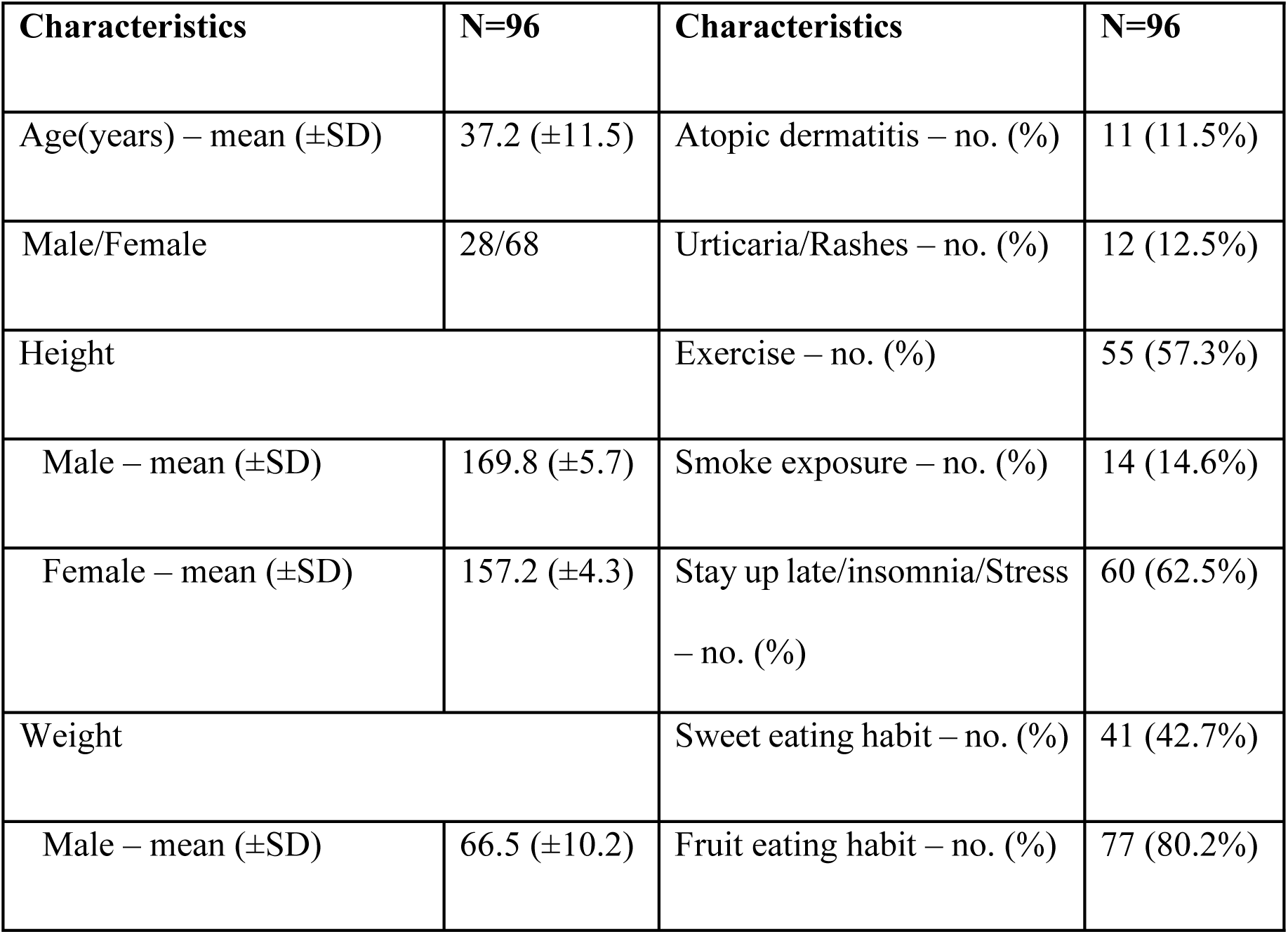
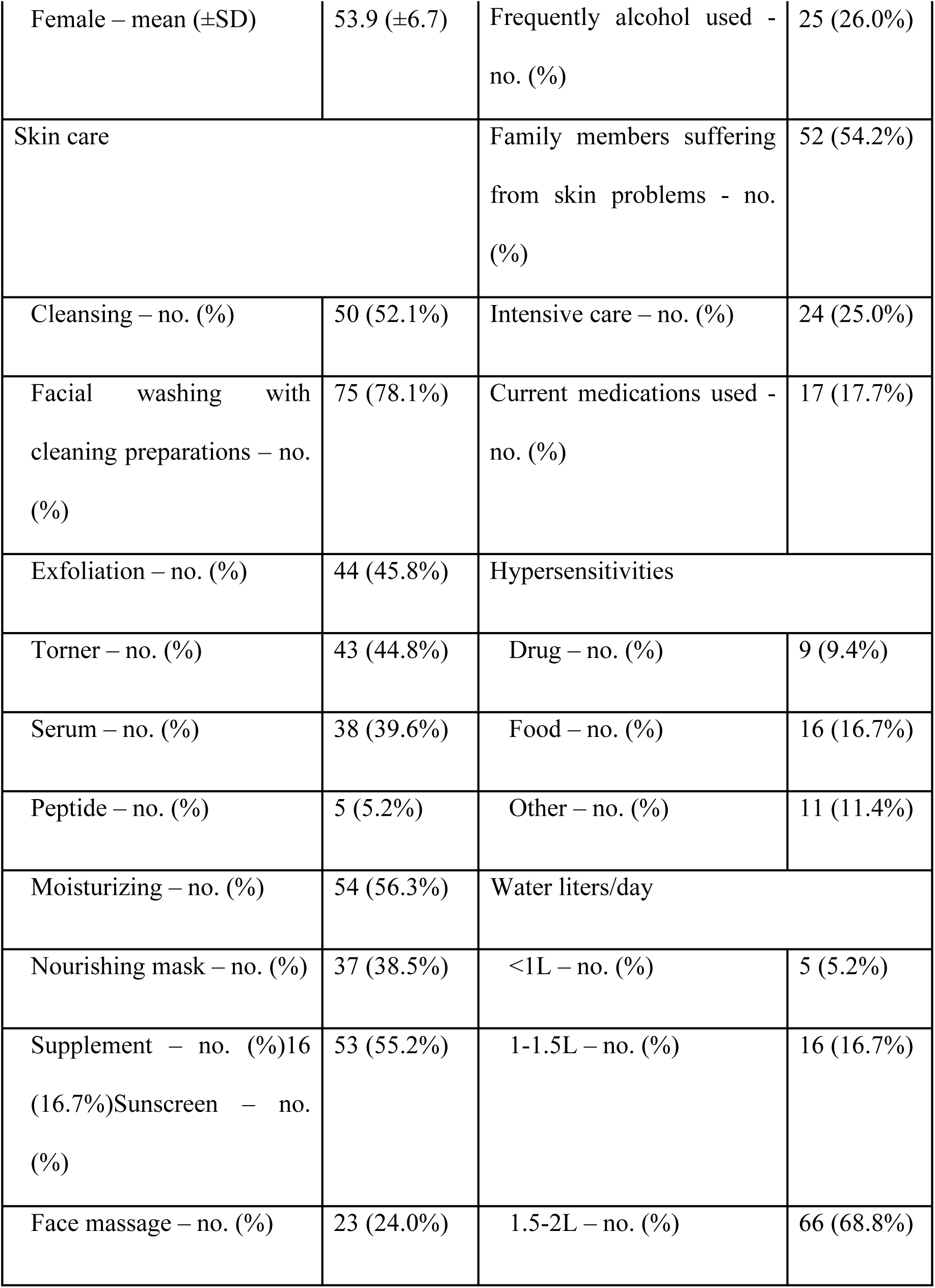
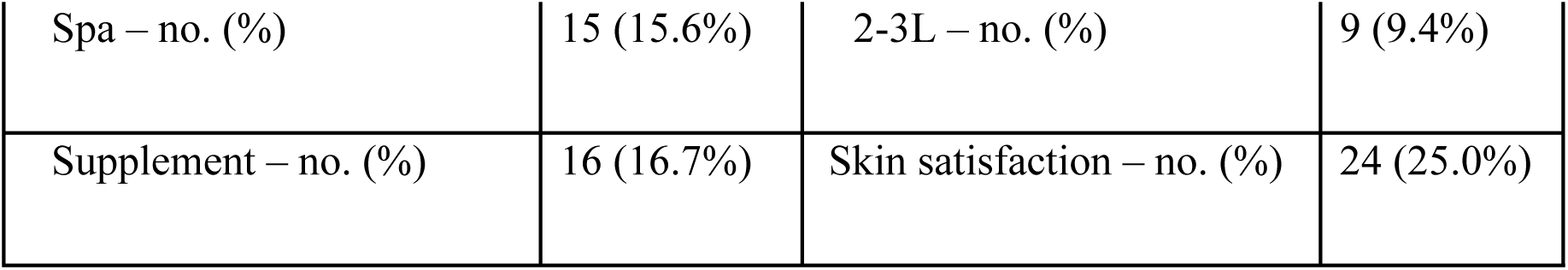
Characteristics of 96 participants involved in the microarray dataset.

| Characteristics | N=96 | Characteristics | N=96 |
| --- | --- | --- | --- |
| Age(years) – mean (±SD) | 37.2 (±11.5) | Atopic dermatitis – no. (%) | 11 (11.5%) |
| Male/Female | 28/68 | Urticaria/Rashes – no. (%) | 12 (12.5%) |
| Height |  | Exercise – no. (%) | 55 (57.3%) |
| Male – mean (±SD) | 169.8 (±5.7) | Smoke exposure – no. (%) | 14 (14.6%) |
| Female – mean (±SD) | 157.2 (±4.3) | Stay up late/insomnia/Stress<br>– no. (%) | 60 (62.5%) |
| Weight |  | Sweet eating habit – no. (%) | 41 (42.7%) |
| Male – mean (±SD) | 66.5 (±10.2) | Fruit eating habit – no. (%) | 77 (80.2%) |
| Female – mean (±SD) | 53.9 (±6.7) | Frequently alcohol used - no. (%) | 25 (26.0%) |
| Skin care |  | Family members suffering from skin problems - no. (%) | 52 (54.2%) |
| Cleansing – no. (%) | 50 (52.1%) | Intensive care – no. (%) | 24 (25.0%) |
| Facial washing with cleaning preparations – no. (%) | 75 (78.1%) | Current medications used - no. (%) | 17 (17.7%) |
| Exfoliation – no. (%) | 44 (45.8%) | Hypersensitivities |  |
| Toner – no. (%) | 43 (44.8%) | Drug – no. (%) | 9 (9.4%) |
| Serum – no. (%) | 38 (39.6%) | Food – no. (%) | 16 (16.7%) |
| Peptide – no. (%) | 5 (5.2%) | Other – no. (%) | 11 (11.4%) |
| Moisturizing – no. (%) | 54 (56.3%) | Water liters/day |  |
| Nourishing mask – no. (%) | 37 (38.5%) | <1L – no. (%) | 5 (5.2%) |
| Supplement – no. (%)<br>16 (16.7%)<br>Sunscreen – no. (%) | 53 (55.2%) | 1-1.5L – no. (%) | 16 (16.7%) |
| Face massage – no. (%) | 23 (24.0%) | 1.5-2L – no. (%) | 66 (68.8%) |
| Spa – no. (%) | 15 (15.6%) | 2-3L – no. (%) | 9 (9.4%) |
| Supplement – no. (%) | 16 (16.7%) | Skin satisfaction – no. (%) | 24 (25.0%) |

### Allele Frequency analysis

We also use Weir and Cokerham’s weighted Fst [14] to compare the genetic structure of the Vietnamese population with other populations. Fst is a parameter of F-statistics developed by Weir and Cokerham to summarize population structure based on the theorem by Wright [16].

### Construction of skin feature landscape for the Vietnamese population in comparison with others

We used the GRS-RAC method to assess individual genetic risk for cutaneous issues for 96 participants in this study and 1008 Vietnamese people from VN1K [10]. Risk alleles (R) and non-risk alleles (N) were counted at each locus. Genotype RR was counted as 2, RN as 1, and NN as 0. Then, the number of risk alleles was summed for each participant from the microarray dataset. In this study, this method was called “Top SNP”.

To detect population-specific risks related to skin traits, we would generate a landscape of skin conditions for Vietnamese and other populations derived from 1KGP3. Based on the range of skin-related SNPs, we divided the population into three intervals according to the quantity of variants carried. Three intervals were named as “Better-than-average”, “Average”, and “Worse-than-average” skin conditions. The process of data curation and cohort categorization is depicted in **Figure 1**.

**Figure 1.**
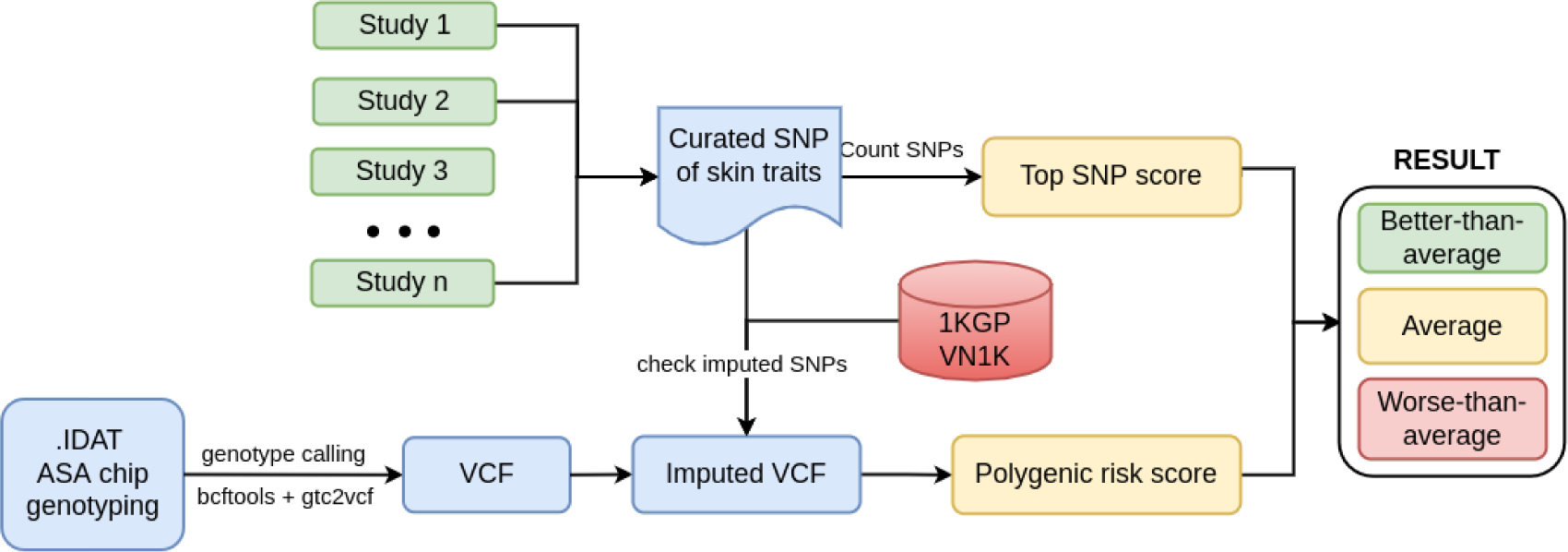
The process of SNP curation and division of VN1K by PRS and Top SNP methods into three equal intervals, equivalent to 3 statuses of “Better-than-average,” “Average,” and “Worse-than-average,” with increased risk of skin-related traits.

### Correlation analysis

The correlations of numerous factors included in the metadata and genetic risk of skin conditions of the microarray dataset, represented by their Top SNP score, were evaluated to find their relationships. The levels of correlations ranged from -1 to 1. The closer R is to 1, the more the correlated variable promotes the latter and vice versa. If R equals 0, there is no correlation between the two variables. The p-value<0.05 showed a significant correlation.

## Results

### The skin variation of Vietnamese compared with other populations

There is a noticeable distinction in allele frequency of variant-associated skin traits between different populations worldwide. We first analyzed the overall population distance using the average Weir and Cokerham’s weighted Fst for 85 aforementioned skin-related SNPs. The genetic profile associated with skin traits of the Vietnamese (VN1K) population was the most similar to the East Asian population. In particular, fixation index values (Fst) between Vietnam and these populations (JPT: Fst=0.036, CHB: Fst=0.031, CHS: Fst=0.027, CDX: Fst=0.025) are far less significant than those between Vietnam and South Asian (SAS), European (EUR), Ad Mixed American (AMR) or African (AFR) (Fst ≥0.09) (**Figure 2**).

**Figure 2.**
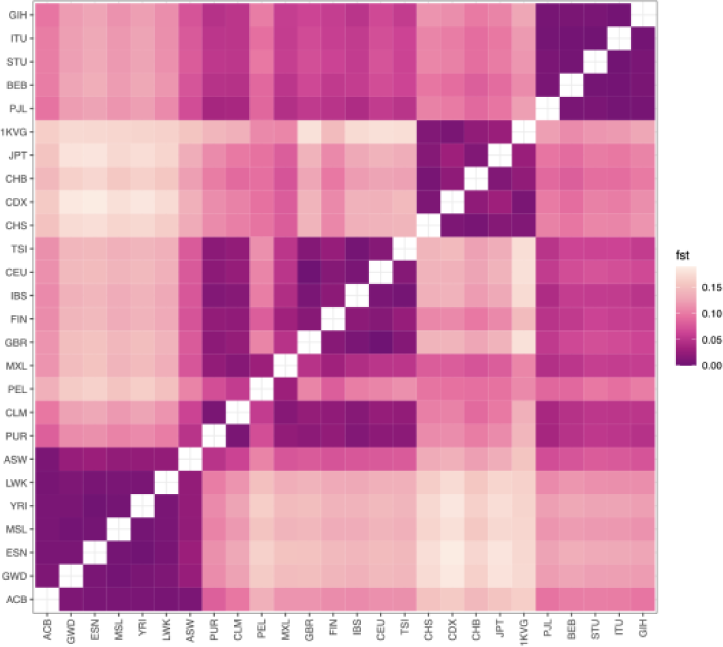
Heatmap displaying Fst score for Vietnamese versus other populations from 1KGP3. 1000 Vietnamese genome project, East Asian (CHS: Han Chinese South; CDX: Chinese Dai in Xishuangbanna, China; CHB: Han Chinese Beijing, China; JPT: Japanese in Tokyo, Japan), South Asian (PJL: Punjabi in Lahore, Pakistan; BEB: Bengali in Bangladesh; STU: Sri Lankan Tamil in the U.K.; ITU: Indian Telugu in the U.K.; GIH: Gujarati Indians in Houston, Texas, USA); European (GBR: British From England and Scotland; FIN: Finnish in Finland; IBS: Iberian Populations in Spain; CEU: Utah residents (CEPH) with Northern and Western European ancestry; TSI: Toscani in Italia), American (PUR: Puerto Rican in Puerto Rico; CLM: Colombian in Medellín, Colombia; PEL: Peruvian in Lima Peru; MXL: Mexican Ancestry in Los Angeles CA USA), African (ACB: African Caribbean in Barbados; GWD: Gambian in Western Division - Mandinka; ESN: Esan in Nigeria; MSL: Mende in Sierra Leone; YRI: Yoruba in Ibadan, Nigeria; LWK: Luhya in Webuye, Kenya; ASW: African Ancestry in SW, USA

### The genomic landscape of skin traits of Vietnamese

Figure 3 shows that the risks vary amongst different skin-related traits in Vietnamese. For example, most Vietnamese people possess an advantageous genetic profile for antioxidant response. In other words, the majority of the Vietnamese population has a remarkable ability to defend their skin against oxidizing agents. In contrast, a higher proportion of the investigated population was shown to be at high risk of moisturizing, collagen degradation, and acne. These conditions would rapidly worsen skin health and accelerate skin aging [17, 18]. Regarding skin-related micronutrient requirements, most Vietnamese people carry a favorable genotype for maintaining adequate serum levels of vitamin D and essential fatty acid (EFA) (e.g., omega-3, omega-6); oppositely, they are likely to be at risk of suffering from deficiency of vitamin C, vitamin B complex and vitamin K.

**Figure 3.**
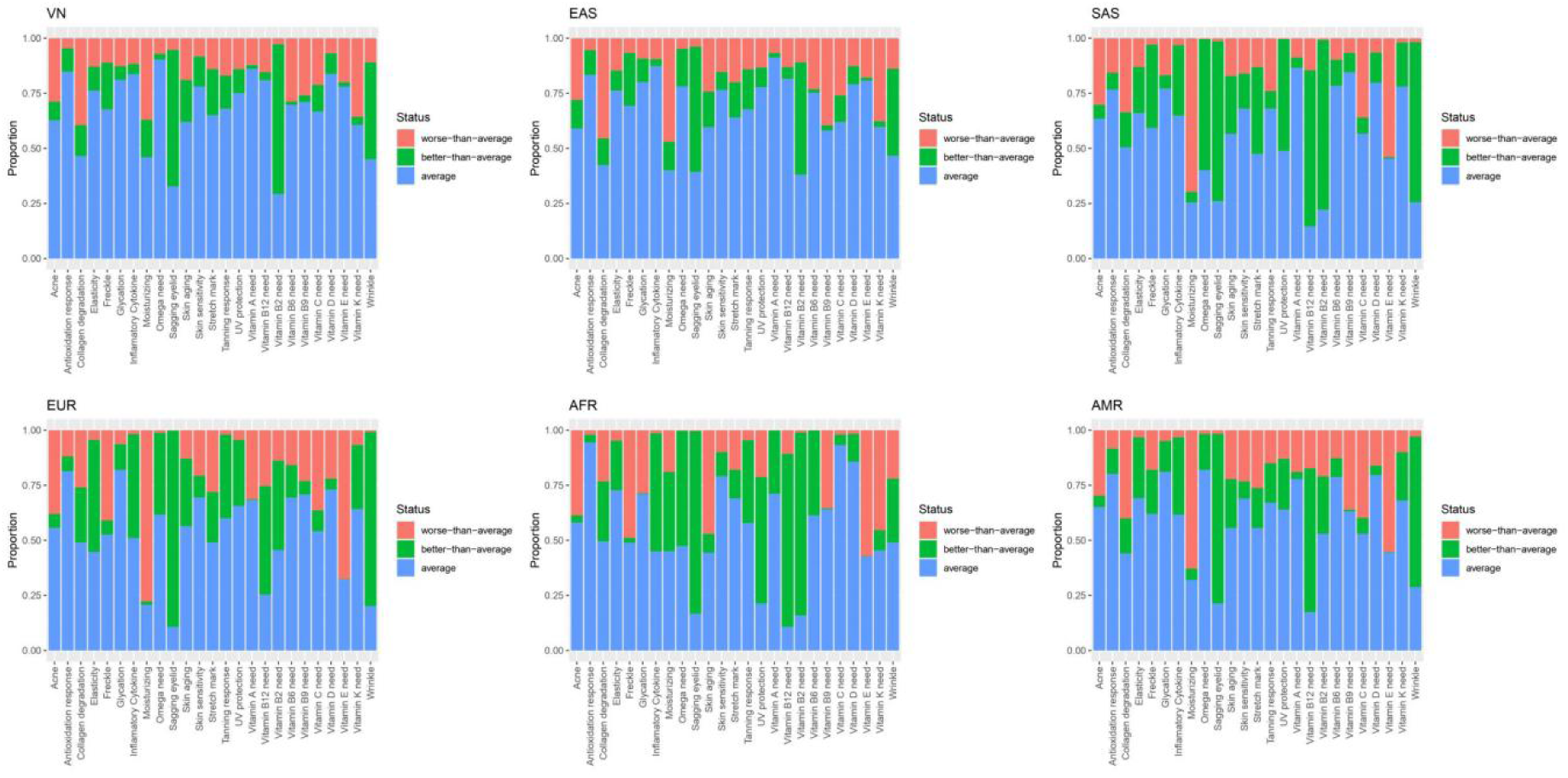
The genomic landscape of skin-related traits in the Vietnamese population and five superpopulation derived from the 1000 genomes project. VN: Vietnamese (VN1K); EAS: East Asian, AFR: African; AMR: American; EUR: European; SAS: South Asian

The genomic landscape of Vietnamese facial skin traits showed the highest similarity level with the East Asian population (EAS). In contrast, there are several differences between the Vietnamese people and other ancestries, such as European, African, and American (**Table S1**). Indeed, European and American genetic profiles have shown to be more susceptible to freckles and moisturizing, whereas African people are at higher risk of wrinkles and glycation than Vietnamese. Four superpopulation from 1KGP3 (except East Asian) are prone to vitamin E deficiency, while more than 75% of the Vietnamese population carry a benign genetic profile for this condition.

### Correlation among skin-related personal characteristics and Top SNP scores

Based on the information retrieved from questionnaire forms and genotyping results of the microarray dataset, a correlation analysis was performed to identify potential relationships between several skin-related traits and genetic profiles (**Table S2**). As expected, we observed the fact that in comparison between the genders, men have higher levels of consuming alcohol (r =-0.51, p = 6.18e-7), and men also do exercise more frequently (r = -0.24, p = 0.026). In contrast, women often skincare and do cosmetic intervention (r = 0.59, p = 5.03e- 09; r = 0.30, p = 0.005, respectively); however, they had lower satisfaction with their skin condition (r = -0.28, p = 0.009). People tend to be interested in their skin health by age, which could be seen through the fact that older people skincare more carefully and frequently (r = 0.26, p = 0.019).

By sharing risk variants, or causality relationships, a few traits show a significant correlation (e.g., collagen degradation and elasticity (r = 0.84, p = 8.0e-4), skin aging and wrinkles (r = 0.45, p =1.50e-5). Aside from these pairs, several skin traits could present in the Vietnamese population, such as collagen degradation often appearing with moisturizing (r=0.35, p = 1.1e- 3), freckles and inflammatory cytokine (r = 0.23, p = 0.03). Moreover, some cutaneous issues tend to come with micronutrient deficiency. It could be seen at the pair of tanning responses and Vitamin B9 level (r = -0.33, p = 2.6e-3) or moisturizing and vitamin A level (r = 0.26, p = 0.02). These findings could suggest their relationship and provide information for constructing the most proper strategy to improve skin health.

## Discussion

The advent of VN1K offers an excellent database for getting more understanding of genomics and their role in various health problems. By using the genetic profile of all participants in VN1K, allele frequency analysis and phenotype landscape have shown a remarkable distinction between populations, suggesting that aside from environmental factors (e.g., climate, living habits, etc.) [19], the genetic profile is an important cause leading to population-specific facial skin issues. These findings could partly explain the different patterns of skin issues in populations. For example, Vietnamese people often deal with collagen degradation and acne, while the significant problems of Europeans are freckles and moisturizing. Therefore, it is crucial to understand the unique skin problems and their causes in each population to properly construct a specific strategy to defend against their high-risk issues.

We identified correlations between several pairs of skin-related traits, which suggested a tendency for their co-occurrences in the Vietnamese population. These phenomena could be attributed to the causality relationship or sharing the related variants between two traits (e.g., collagen degradation and elasticity, or skin aging and wrinkle); however, in cases of the remaining pairs (e.g., collagen degradation and moisturizing, moisturizing and Vitamin A level, or freckle and inflammatory cytokine, etc.) and might suggest the potential associations, particularly in Vietnamese individuals.

## Conclusion

This is the first study on the Vietnamese population to investigate the genetic features which play an essential role in skin health. However, further functional characterizations of the investigated genes are warranted to elucidate their contribution to skin-related traits. By examining several skin-associated genetic variants in the Vietnamese population, this study could improve inadequate skin-related genetic diversity in the currently published database.

## Abbreviations

VN1K: 1000 Vietnamese Genomes Project
1KGP3: 1000 Genomes Project Phase 3
AFR: African
AMR: Ad Mixed American
ASA: Infinium Asian Screening Array- 24 v1.0 BeadChip
EAS: East Asian
EFA: Essential fatty acid
EUR: European
GWAS: Genome-wide association studies
LD: Linkage disequilibrium
NGS: Next-generation sequencing
PRS: Polygenic Risk Score
SAS: South Asian
SNP: Single Nucleotide Polymorphism
WGS: Whole Genome Sequencing.

## Acknowledgments

We would like to thank our colleagues at Vingroup Big Data Institute, GeneStory JSC, and Vinmec Healthcare System, especially Dr. Nguyen Thuy Duong for her feedback on the questionnaire, Ms. Tran Thi Ha Trang for her contribution to VN1K.

## Authorship contribution statement

Tham Hong Hoang: Conceptualization, Roles/Writing - original draft, Writing - review & editing, Formal analysis. Duc Minh Vu: Data curation, Investigation, Funding acquisition, Project administration. Giang Minh Vu: Roles/Writing - original draft, Formal analysis. Thien Khac Nguyen: Writing - review & editing, Data curation. Nguyet Minh Do: Roles/Writing - initial draft, Data curation. Vinh Chi Duong: Data curation. Thang Luong Pham: Investigation, Methodology. Mai Hoang Tran: Investigation, Methodology. Ly Thi Khanh Nguyen: Methodology. Han Thi Tuong Han: Data curation. Thuy Thu Can: Data curation. Thai Hong Pham: Investigation, Methodology. Tho Duc Pham: Investigation, Methodology. Thanh Hong Nguyen: Investigation, Methodology. Huy Phuoc Do: Investigation, Methodology. Nam S. Vo: Conceptualization, Funding acquisition, Project administration. Xuan-Hung Nguyen: Conceptualization, Funding acquisition, Project administration, Writing - review & editing.

## Ethics declaration

This study was approved by the Ethics Committee in Biomedical Research of Vinmec Healthcare System 03/2022/CN-HDDD VMEC. Personal information is entirely confidential. In the VN1K study, subjects provided informed consent, and the Vinmec International Hospital Institutional Review Board approved the study with number 543/2019/QƉ-VMEC. All methods were carried out in accordance with the relevant guidelines and regulations (e.g., Helsinki Declaration).

## Figure captions

**Figure S1.**
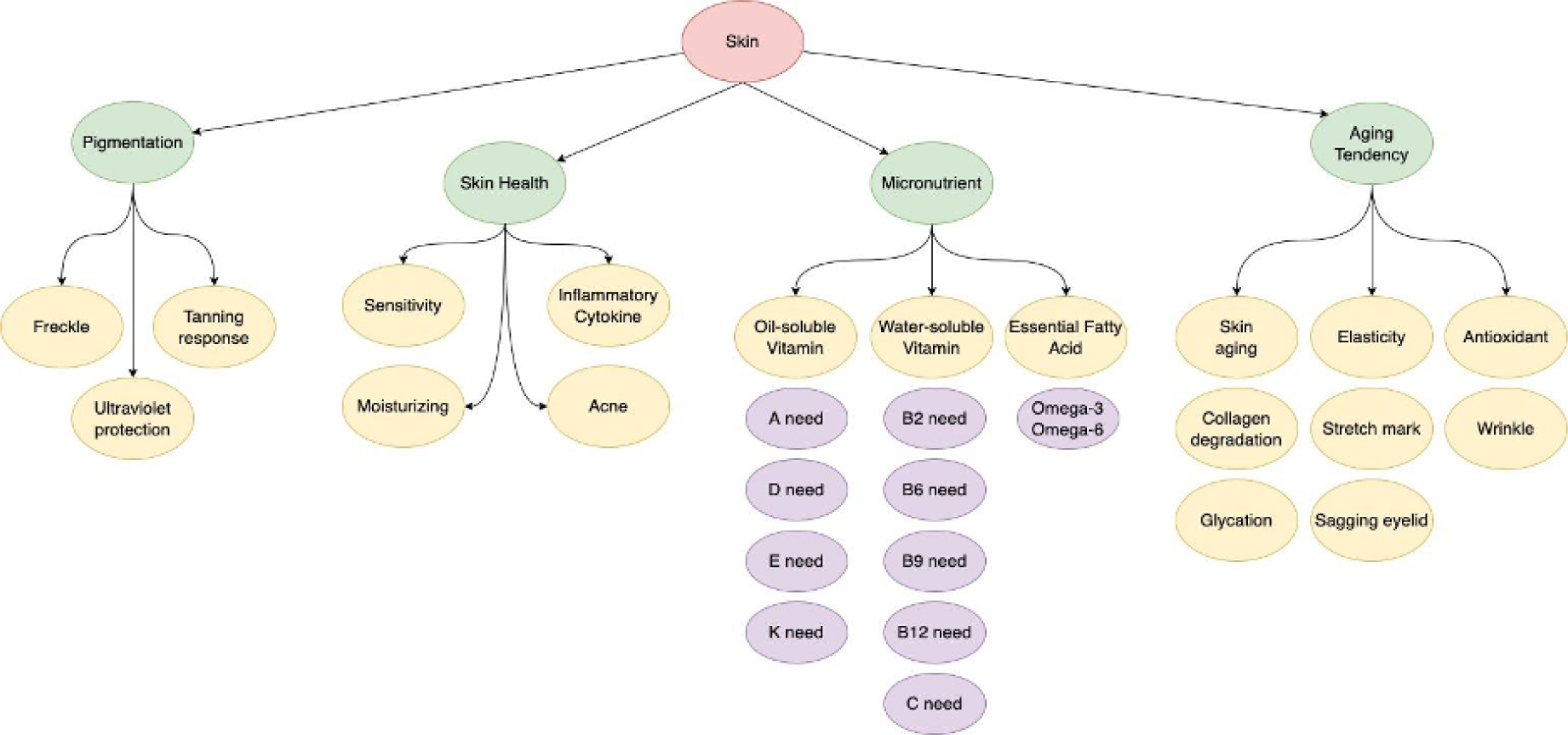
The structure of 25 skin-related traits illustrates relationships in four groups of Pigmentation, Dermatitis, Nutrition, and Aging.

**Table S1.**
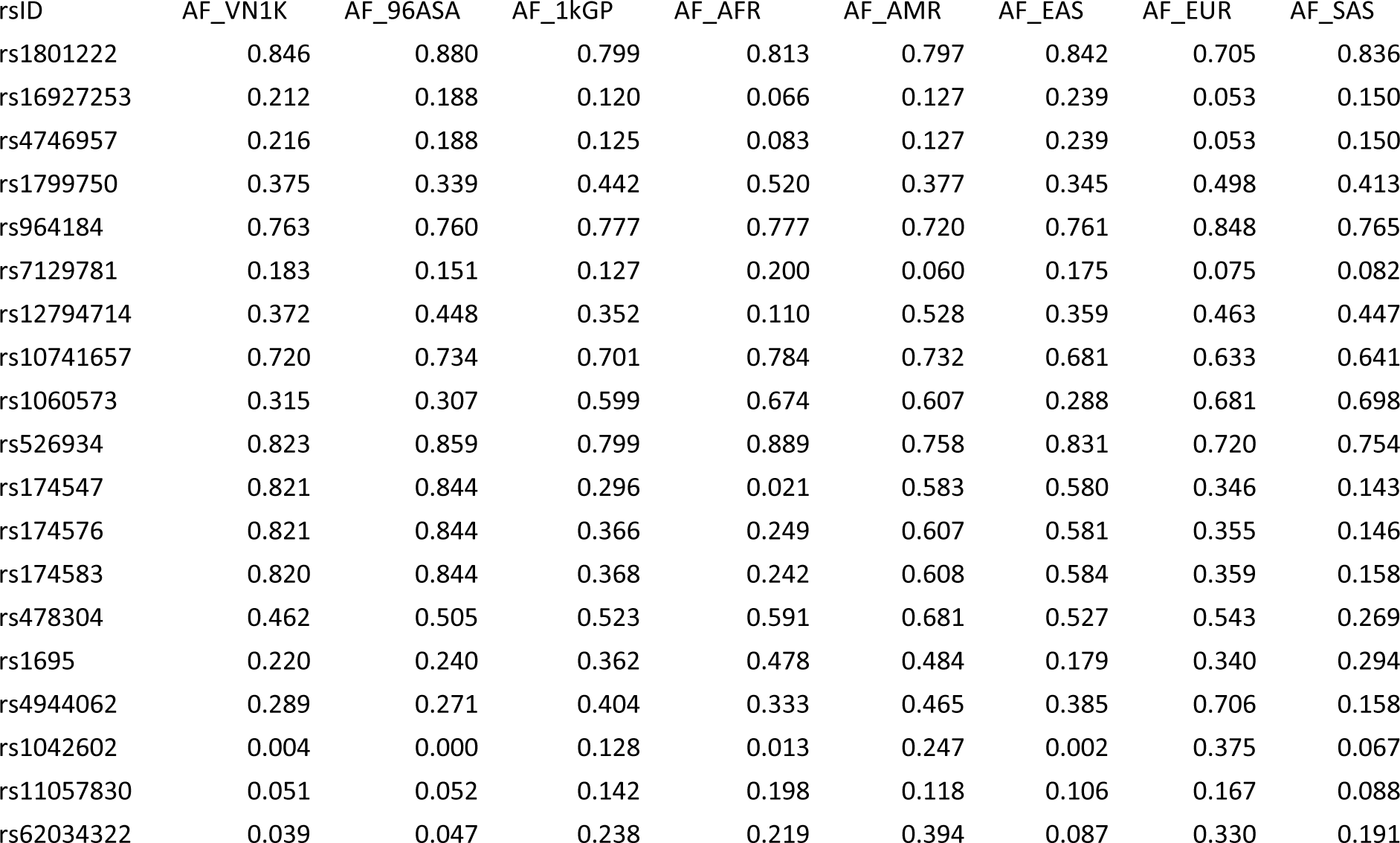
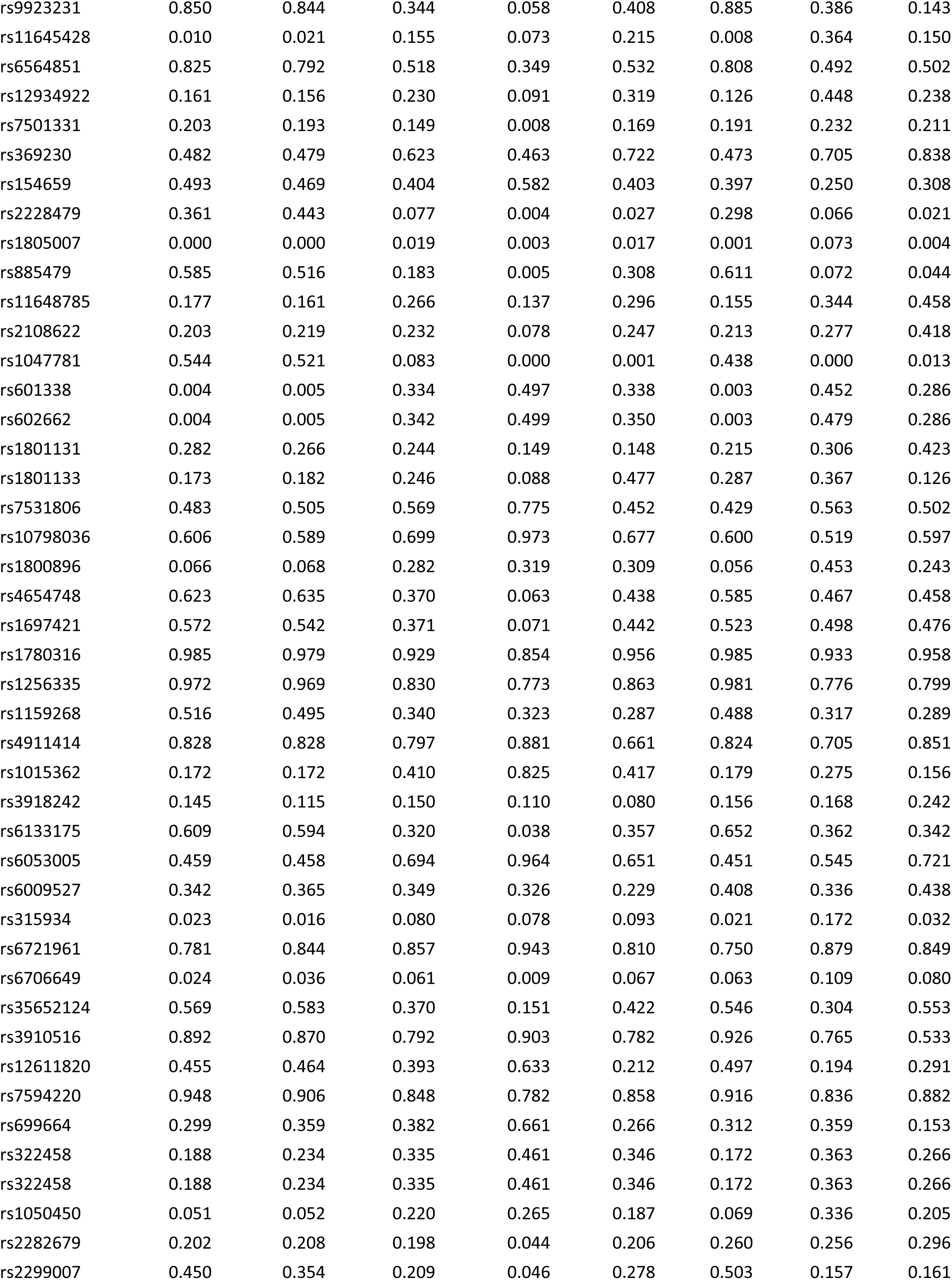
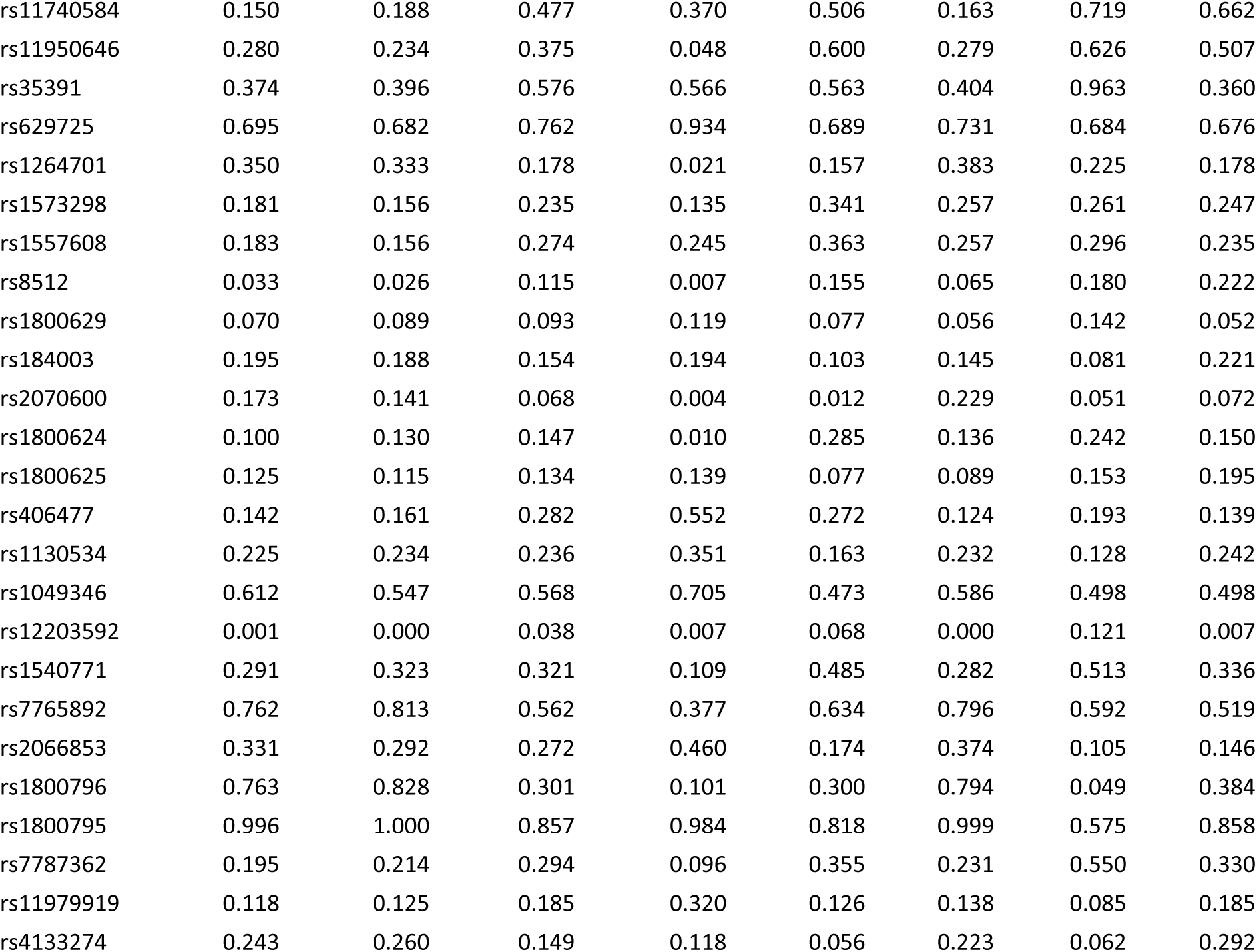
The allele frequency of skin-related SNPs.

## Reference

[1] R.J. Hay, N.E. Johns, H.C. Williams, I.W. Bolliger, R.P. Dellavalle, D.J. Margolis, et al., The global burden of skin disease in 2010: an analysis of the prevalence and impact of skin conditions, J Invest Dermatol 134(6) (2014) 1527–1534.

[2] K. Park, Role of micronutrients in skin health and function, Biomol Ther (Seoul) 23(3) (2015) 207–17.

[3] C. Endo, T.A. Johnson, R. Morino, K. Nakazono, S. Kamitsuji, M. Akita, et al., Genome-wide association study in Japanese females identifies fifteen novel skin-related trait associations, Sci Rep 8(1) (2018) 8974.

[4] M. Jonnalagadda, M.A. Faizan, S. Ozarkar, R. Ashma, S. Kulkarni, H.L. Norton, et al., A Genome-Wide Association Study of Skin and Iris Pigmentation among Individuals of South Asian Ancestry, Genome Biol Evol 11(4) (2019) 1066–1076.

[5] P.R. Loh, G. Kichaev, S. Gazal, A.P. Schoech, A.L. Price, Mixed-model association for biobank- scale datasets, Nat Genet 50(7) (2018) 906–908.

[6] J.Y. Seo, S.W. You, J.G. Shin, Y. Kim, S.G. Park, H.H. Won, et al., GWAS Identifies Multiple Genetic Loci for Skin Color in Korean Women, J Invest Dermatol 142(4) (2022) 1077–1084.

[7] R.A. Sturm, D.L. Duffy, Human pigmentation genes under environmental selection, Genome Biol 13(9) (2012) 248.

[8] Y. Liu, W. Gao, C. Koellmann, S. Le Clerc, A. Hüls, B. Li, et al., Genome-wide scan identified genetic variants associated with skin aging in a Chinese female population, Journal of Dermatological Science 96(1) (2019) 42–49.

[9] S.W. Choi, T.S. Mak, P.F. O’Reilly, Tutorial: a guide to performing polygenic risk score analyses, Nat Protoc 15(9) (2020) 2759–2772.

[10] R. Na, D. Ye, J. Qi, F. Liu, X. Lin, B.T. Helfand, et al., Race-specific genetic risk score is more accurate than nonrace-specific genetic risk score for predicting prostate cancer and high-grade diseases, Asian J Androl 18(4) (2016) 525–9.

[11] Infinium asian screening array beadchip data sheet.

[12] O. Delaneau, J.F. Zagury, M.R. Robinson, J.L. Marchini, E.T. Dermitzakis, Accurate, scalable and integrative haplotype estimation, Nat Commun 10(1) (2019) 5436.

[13] S. Das, L. Forer, S. Schönherr, C. Sidore, A.E. Locke, A. Kwong, et al., Next-generation genotype imputation service and methods, Nat Genet 48(10) (2016) 1284–1287.

[14] B.S. Weir, C.C. Cockerham, ESTIMATING F-STATISTICS FOR THE ANALYSIS OF POPULATION STRUCTURE, Evolution 38(6) (1984) 1358–1370.

[15] S. Wright, The Interpretation of Population Structure by F-Statistics with Special Regard to Systems of Mating, Evolution 19(3) (1965) 395–420.

[16] H. Zhu, X. Zhou, Statistical methods for SNP heritability estimation and partition: A review, Comput Struct Biotechnol J 18 (2020) 1557–1568.

[17] K. Nolan, E. Marmur, Moisturizers: reality and the skin benefits, Dermatol Ther 25(3) (2012) 229–33.

[18] C. Zip, The Role of Skin Care in Optimizing Treatment of Acne and Rosacea, Skin Therapy Lett 22(3) (2017) 5–7.

[19] J. English, R. Dawe, J. Ferguson, Environmental effects and skin disease, British Medical Bulletin 68(1) (2003) 129–142.

[20] L.C. Jacobs, M.A. Hamer, D.A. Gunn, J. Deelen, J.S. Lall, D. van Heemst, et al., A Genome- Wide Association Study Identifies the Skin Color Genes IRF4, MC1R, ASIP, and BNC2 Influencing Facial Pigmented Spots, J Invest Dermatol 135(7) (2015) 1735-1742.

[21] M.H. Law, S.E. Medland, G. Zhu, S. Yazar, A. Viñuela, L. Wallace, et al., Genome-Wide Association Shows that Pigmentation Genes Play a Role in Skin Aging, J Invest Dermatol 137(9) (2017) 1887–1894.

[22] M.A. Farage, Y. Jiang, J.P. Tiesman, P. Fontanillas, R. Osborne, Genome-Wide Association Study Identifies Loci Associated with Sensitive Skin, Cosmetics 7(2) (2020) 49.

[23] M.E. Breitbach, S. Greenspan, N.M. Resnick, S. Perera, A.U. Gurkar, D. Absher, et al., Exonic Variants in Aging-Related Genes Are Predictive of Phenotypic Aging Status, Front Genet 10 (2019) 1277.

[24] K. Adhikari, J. Mendoza-Revilla, A. Sohail, M. Fuentes-Guajardo, J. Lampert, J.C. Chacón-Duque, et al., A GWAS in Latin Americans highlights the convergent evolution of lighter skin pigmentation in Eurasia, Nat Commun 10(1) (2019) 358.

[25] V. Roberts, B. Main, N.J. Timpson, S. Haworth, Genome-Wide Association Study Identifies Genetic Associations with Perceived Age, J Invest Dermatol 140(12) (2020) 2380–2385.

[26] K. Ganguly, T. Saha, A. Saha, T. Dutta, S. Banerjee, D. Sengupta, et al., Meta-analysis and prioritization of human skin pigmentation-associated GWAS-SNPs using ENCODE data-based web- tools, Arch Dermatol Res 311(3) (2019) 163–171.

[27] I. Galván-Femenía, M. Obón-Santacana, D. Piñeyro, M. Guindo-Martinez, X. Duran, A. Carreras, et al., Multitrait genome association analysis identifies new susceptibility genes for human anthropometric variation in the GCAT cohort, J Med Genet 55(11) (2018) 765–778.

